# Gut transcriptome reveals differential gene expression and enriched pathways linked to immune activation in response to weaning in pigs

**DOI:** 10.1101/2021.11.30.470420

**Authors:** M. Le Bon, S. Tötemeyer, R. D. Emes, K. H. Mellits

**Affiliations:** School of Biosciences, Division of Microbiology, Brewing and Biotechnology, University of Nottingham, Loughborough, LE12 5RD, UK; School of Animal, Rural and Environmental Sciences, Nottingham Trent University, Southwell, NG25 0QF, UK; School of Veterinary Medicine and Science, University of Nottingham. LE12 5RD, UK; Advanced Data Analysis Centre, University of Nottingham. LE12 5RD, UK

**Author notes:** Corresponding author: M. Le Bon.

**Keywords:** Pig, Weaning, RNA-sequencing, Transcriptomic, Gut, Immune response

## Abstract

Weaning represents one of the most critical periods in pig production associated with increase in disease risk, reduction in performance and economic loss. Physiological changes faced by piglets during the weaning period have been well characterised, however little is currently known about the underlying molecular pathways involved in these processes. As pig meat remains one of the most consumed sources of protein worldwide, understanding how these changes are mediated is critical to improve pig production and consequently sustainable food production globally. In this study, we evaluated the effect of weaning on transcriptomic changes in the colon of healthy piglets over time using an RNA-sequencing approach.

The findings revealed a complex and coordinated response to weaning with the majority of genes found to be rapidly differentially expressed within one day post weaning. Multiple genes and pathways affected by weaning in the colon were associated with immune regulation, cell signalling and bacterial defence. NOD-like receptors, Toll-like receptor and JAK-STAT signalling pathways were amongst the pathways significantly enriched. Immune activation was evidenced by the enrichment of pathways involved in interferon response, cytokines interactions, oxidoreductase activities and response to microbial invasion. Biosynthesis of amino acids, in particular arginine, was also amongst the most enriched KEGG pathways in weaned pigs, reinforcing the critical role of arginine in gut homeostasis under stress conditions.

Overall, transcriptomic and physiological results suggest that pigs going through the weaning transition undergo a transient period of inflammatory state with a temporary breakdown of barrier functions in the gut. These findings could provide valuable tools to monitor host response post weaning, and may be of particular relevance for the investigation and development of intervention strategies aimed to reduce antibiotic use and improve pig health and performance.

## Introduction

Livestock production is expected to produce more food than ever before. As the expanding world population is getting wealthier, the demand for safe and secure animal protein is increasing (Henchion et al., 2017). The challenge is to meet this demand in ways that are environmentally, socially and economically sustainable. Together with poultry, pork is one of the fastest growing livestock sectors and also one of the most consumed meats world-wide (FAO, 2019). Pig production is also widely recognised as one of the most efficient in term of carbon footprint and climate change potential compared to other animal protein source (Macleod et al., 2013). Ensuring optimal production is critical to ensure animals can fulfil their genetic potential and contribute to sustainable food source.

As a result of abrupt dietary, social stresses, and environmental changes, weaning is recognised as the most critical period in modern pig production associated with increase in disease risks, reduction in performance and welfare leading to significant economic loss (Gresse et al., 2017; Nowland et al., 2019). At weaning, the pig gastrointestinal tract (GIT) undergoes rapid changes in size, protein turnover rates, microbiome composition, and detrimental alterations in digestive and barrier functions (Pluske et al., 2018). Although the physiological changes faced by piglets over weaning have been well characterised, little is known about the underlying genes and pathways involved in these processes. Understanding how these changes are regulated or mediated is critical to improve pig production and consequently sustainable food production globally.

Furthermore, due to similarities in anatomy and physiology, the pig is widely recognised and utilised as a translational animal model to study human gastrointestinal diseases and to understand biological pathways related to mucosal function, development and nutritional regulation (Roura et al., 2016; Sciascia et al., 2016; Zhang et al., 2013). Previous studies have highlighted the importance of improving the knowledge on molecular mechanism responsible for phenotypic differences especially at an early ages with the dual purpose of improving production and providing adequate models for human studies (Ayuso et al., 2015).

Recent advances in sequencing technologies now provides novel opportunities to comprehensively explore the complex gut ecosystem of humans and animals. RNA-sequencing (RNA-Seq) is a powerful high-throughput approach to profile gene expression that, in contrast with microarray-based technologies, allows for the characterisation and quantification of both known and unknown transcripts (Mach et al., 2014). Fundamental understanding of the host response to stress and its environment is paramount for the optimisation and development of application to improve pig health, productivity and welfare. To date, RNA-Seq has been used to study production traits of livestock animals but transcriptomic study in pigs using this technology is relatively scarce and have mainly focused on disease response or regulation mechanism of fat deposition and muscle development to evaluate growth and meat quality between pig genotypes (Piórkowska et al., 2018; Xu et al., 2019).

Much attention and focus has been given to the gut microbiome and its taxonomic and metabolic changes through pig development and weaning (Bian et al., 2016; Frese et al., 2015; Guevarra et al., 2018a) but we are still lacking understanding about the host gene expression change in response to weaning. The current study aims to investigate the transcriptomic changes in the pig gut through weaning over time.

## Material and methods

### Animals and experimental design

All animals were treated in accordance with the University of Nottingham ethical guidelines and codes of practice applying to care and management of animals. Twenty-four litters (Landrace x Large White) over three batches were used in the study and housed at the School of Biosciences, Sutton Bonington Campus, University of Nottingham, UK. For prevention of iron deficiency and coccidiosis, all piglets received a 1 ml IM injection of Gleptosil (Alstoe Ltd, York, UK) 24 h after birth, and 0.7 ml of Baycox (Bayer, Newbury, UK) orally 3 d after birth. At 21 days of age, 6 weight-matched piglets per litter were randomly allocated by random selection of coded balls to treatment (baseline average weight at day 19: 6.77 ± 0.189 kg). Piglets allocated to the weaned treatment were separated from their dam, moved and mixed with non-littermates in pens of 4 individuals and received ad lib commercial diet (wheat, whey powder and soya based) containing: 21.25% protein, 7.5% fat, 2.00% fibre, 5% ash, 1.70% lysine, 13.80% moisture). Weaned piglets did not receive creep feed supplementation before weaning. Piglets allocated to the unweaned treatment as control remained with their dam and littermate up to 35 days of age with access to creep feed from day 25d of age (same commercial diet as above). No antibiotic or anthelmintic treatment were used during the trial. At day 1, 4 and 14 post-weaning: one weaned and one unweaned piglet from each litter were weighed euthanised by intraperitoneal injection of Dolethal (1 ml.kg-1 body weight; 20% w/v Pentobarbitone Sodium, Vétoquinol, Buckingham, UK). At slaughter, body lesion was scored for each pig on a 3 point scale with 1 for no lesion, 2 for moderate scratches on back, flank, head, ear and tail, and 3 for intense, deep or bleeding scratches on back, flank, head, ear and tail.

### Intestinal measurements

All sample processing and analysis was blinded using randomly generated numerical codes. Tissue samples from the 0.5 section of the colon were immediately collected post-slaughter, rinsed in sterile buffered saline solution and preserved in RNA later (Ambion, CA USA) at 4°C for 24-48h to allow tissue penetration then stored at −80°C. Tissue samples of the 0.5 small intestine (as proportions) along from the gastric pylorus to the ileocecal valve were fixed in Bouin’s solution, embedded in paraffin and cut in 5 µm transverse sections. Histological section were stained with H&E for histometric measurement of villus length and crypt depth, with Periodic Acid Schiff stain for goblet cell counts (Matsuo et al., 1997) and with Toluidine Blue for quantification of mast cells (Moeser et al., 2007).

Secretory IgA (sIgA) was measured in ileal flushes using the methods previously described by Lessard et al. (2009). At slaughter, a 20 cm segment of ileum taken upstream from the cecum was flushed using 5 ml of sterile PBS, and centrifuged for 10 min at 500 g. The supernatant was collected and stored at −80°C. Secretory IgA was measured in duplicate using a sandwich Porcine IgA ELISA Quantitation Kit (Bethyl Laboratories, TX, USA) according to the manufacturer protocol.

### Data and Bioinformatics analysis

Statistical analysis was performed in IBM SPSS v24 to determine the effect of weaning treatment at different time point treatment on pig weight, intestinal and blood measurement using linear mixed model analysis.

Gene expression cluster analysis was performed and revealed that sex was the only factor that showed a grouping effect. Therefore sex was included in the model as a confounder for the gene expression analysis.

Sequence alignment and read quantification was performed using the pseudo-alignment-based tool Kallisto v0.43. (Bray et al., 2016). Differential expression was determined using the Wald test in Sleuth v0.28.1 (Pimentel et al., 2017) with sex as a confounder in the model. Transcripts with a false detection rate corrected p-value < 0.05 and a log2 fold change (log2FC) greater than 1 and less than minus 1 were considered to be differentially expressed, unless otherwise stated. False detection rate correction was performed using the Benjamini-Hochberg method (Benjamini and Hochberg, 1995).

## Results

### Phenotypic data

All piglets were found in good health during the trial and displayed no clinical signs of disease or scour. As expected, weaning caused a number physiological changes which were found to be time-dependant. Significant differences were observed in pig weight, blood or plasma measurements, body lesion scores and intestinal measurement (Table 1). These preliminary results also indicate the significant effect of time on almost all variables measured between day 1 and day 14 post weaning. This highlights that the pigs are undergoing rapid period of development at this age and the importance of designing studies using age-matched controls when evaluating the effect of weaning in pigs as opposed to using pre-weaning values as controls.

**Table 1:**
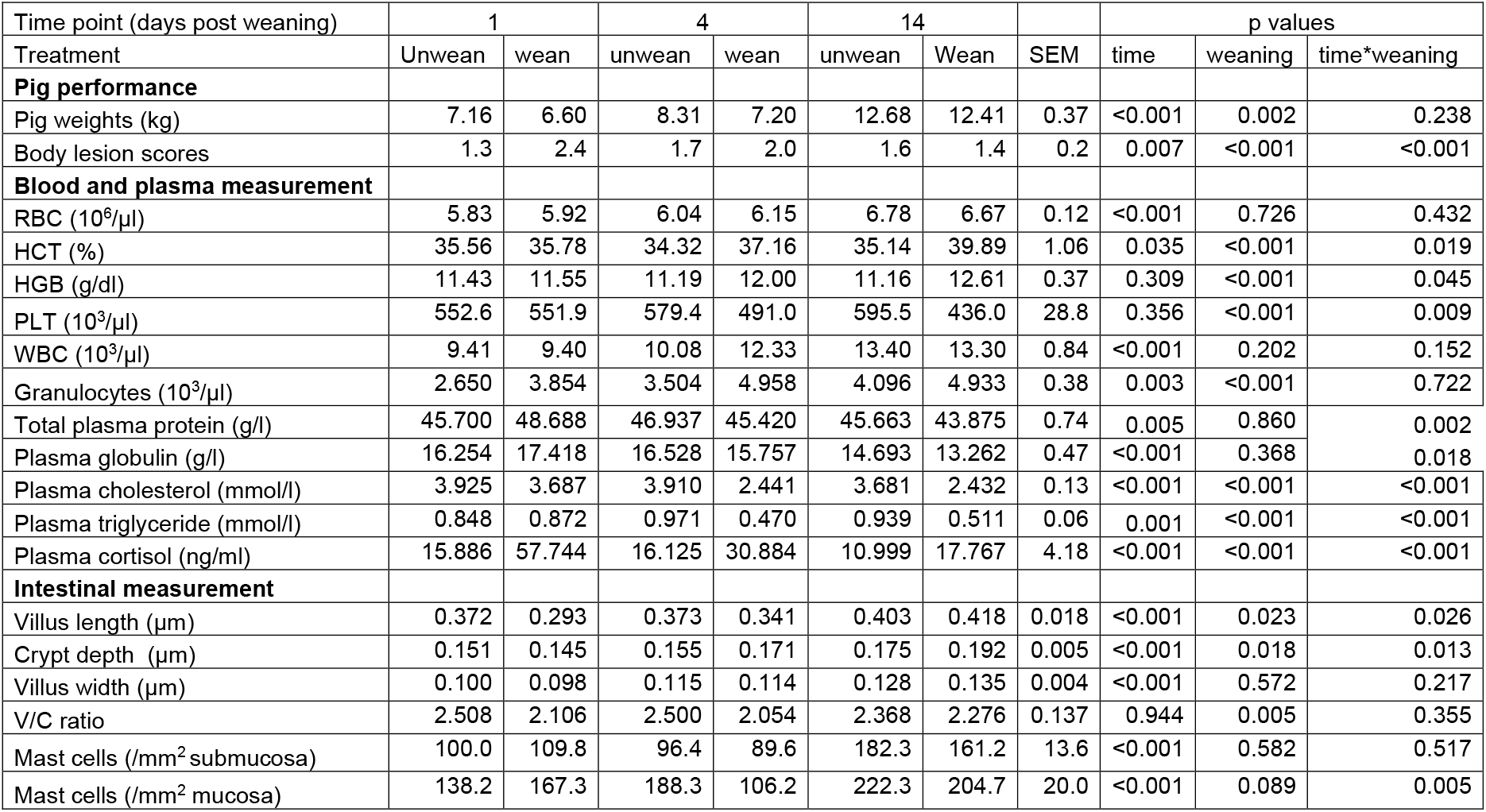
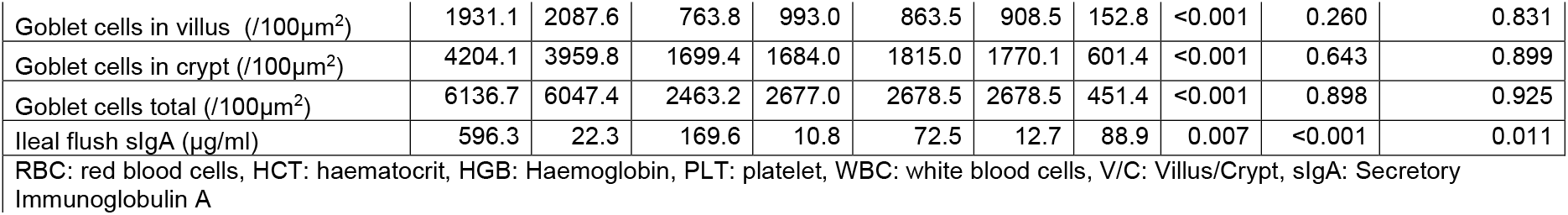
Phenotypic measurements between weaned and unweaned pigs at 1, 4 and 14 days post weaning. Data shown are means and the pooled SEM for weaned and unweaned pigs for each time points.

Plasma analysis revealed that cortisol levels remained stable over time in the unweaned group but increased almost 4-fold at day 1 and 2-fold at day 4 post weaning in the weaned pigs indicating an activation of the HPA axis under weaning stress (Martínez-Miró et al., 2016). Interestingly, while most parameters measured in this study showed a rapid spike followed by a progressive return to the unweaned level by day 14, cortisol level still remained significantly higher at 14 days post weaning although to a lesser magnitude.

Haematology and biochemistry profiles are used as indicators of health status to evaluate the metabolic, nutritional and energy state of the pig. In blood, circulating granulocyte levels, haemoglobin and haematocrit were significantly increased in weaned pigs, while platelet counts decreased. In plasma, cholesterol, triglycerides and globulin levels decreased at day 4 and day 14 in the weaned group compared to unweaned pigs.

Measurement of intestinal architecture were also affected by weaning as previously reported, with significant reduction in jejunal villus height and villus/crypt ratio. However, villus width remains unaffected by weaning and crypt depth was increased. Goblet cell numbers in crypt and in villus remained unaffected in this study, however there was a significant decrease in mucosal mast cells at day 4 with an overall statistical trend for weaned pigs to show reduced mast cell count compared to unweaned controls.

Ileal sIgA was also greatly reduced at all time points post weaning, suggesting a decrease in immune protection from maternal milk. IgA plays an important role in the protection of mucosal surfaces against pathogens and is the principal immunoglobulin secreted in sow’s milk (40% of the total whey protein) (Klobasa et al., 1987). sIgA is considered the first line of defence and one of the most important factor for piglet growth and survival (Salmon et al., 2009). At weaning, removal of the piglets from the sow causes dramatic drop in gut IgA levels as observed in the current study and by others (Lessard et al., 2009), increasing the immune vulnerability of piglets post weaning.

### Transcriptomic analysis

RNAseq (HiSeq) was used to identify deferentially expressed transcripts (DET) between weaned and unweaned pigs in colonic tissue. An average of 52.3 million trimmed paired reads were obtained for each sample. Reads mapped as pairs (83.0-85.3%) to the porcine genome ref sequence (sus scrofa 10.2). The Wald test was used on TPM to establish the total number of DET between weaned and unweaned pigs for each time point and identified a total of 239 transcripts at q value ≤ 0.1 and FC ≥ 2. The volcano plots visually represent significant DET for each time point and show an even distribution between up- and down-regulated genes (Figure 1).

**Figure 1:**
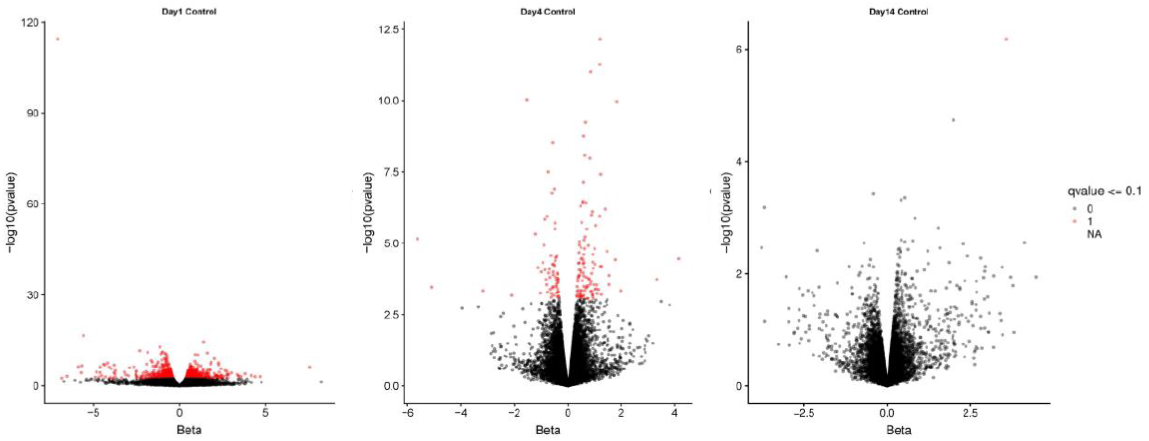
Volcano plots of differentially expressed transcripts between weaned and unweaned pigs at 1, 4 and 14 days post weaning. Red points indicates genes significant at Qval ≤ 0.1.

The majority of transcripts 171 (71.6%) were found to be differentially expressed after 1 day post weaning. After 4 days and 14 days post weaning only 67 (28.0%) and 1 (0.4%) genes were differentially expressed between weaned and unweaned pigs, respectively (Figure 2). Among DET with the cut off values of Qval ≤ 0.1 & FC ≥ 2, two transcripts were identified in common between day 1 and day 4 post weaning: ENSSSCT00000014128 and ENSSSCT00000017308 which are of unknown function (Figure 3). They were no DET common at all three time point or shared between day 4 and 14 or day 1 and 14 post weaning. Transcription according to time point and weaning treatment are shown in PCA plots and revealed distinct clusters between weaned and unweaned pigs across time points (Figure 4).

**Figure 2:**
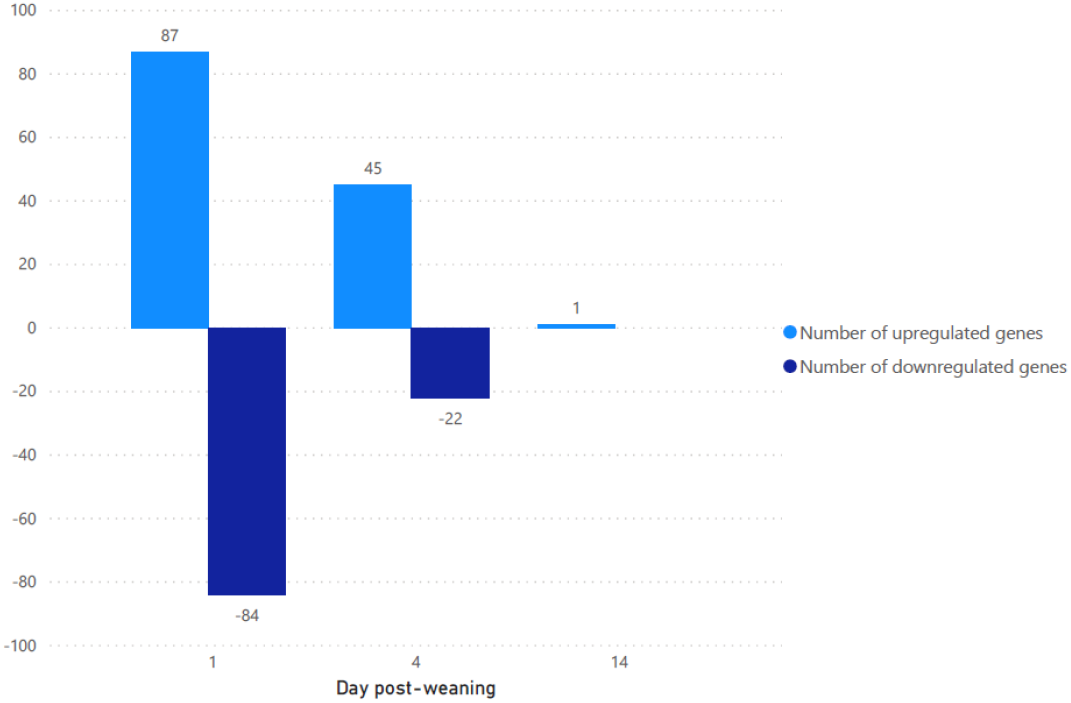
Number of up and downregulated transcripts between weaned and unweaned pigs at 1, 4 and 14 days post weaning (Qval ≤ 0.1 & FC ≥ 2).

**Figure 3:**
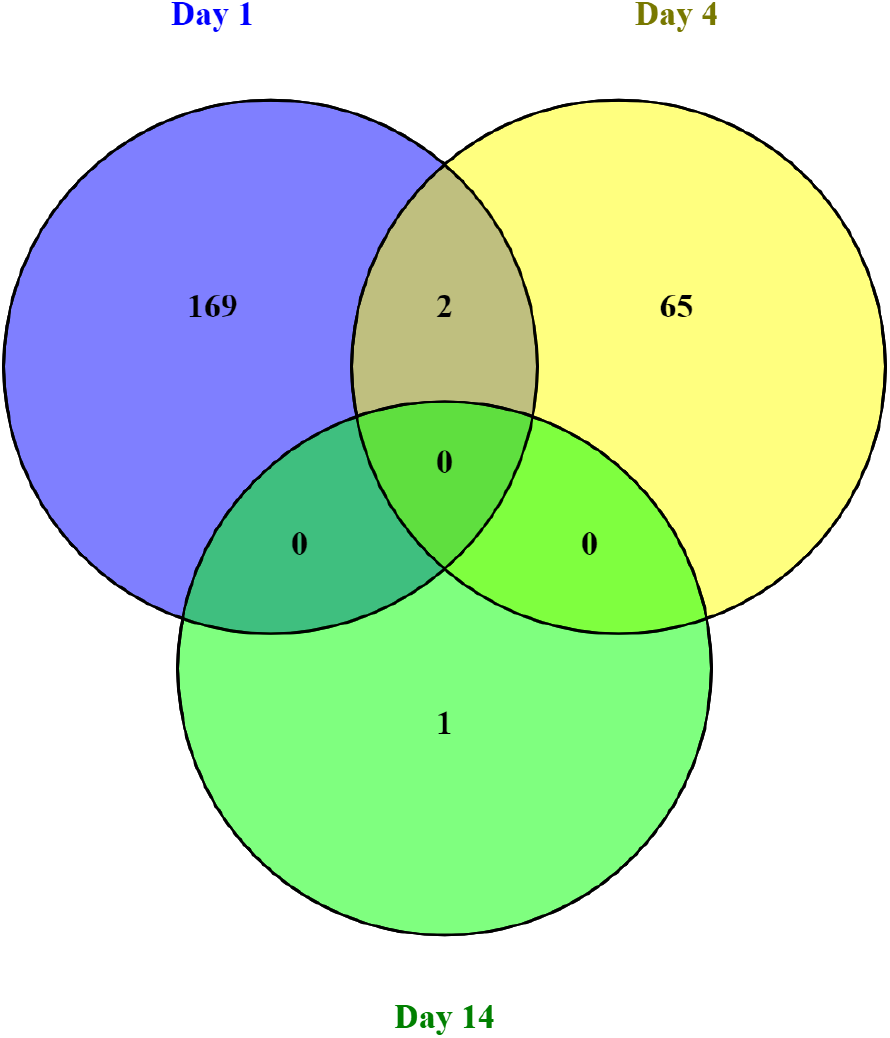
Venn diagram of the number of differentially expressed transcripts between weaned and unweaned pigs for each time point (Qval ≤ 0.1 & FC ≥ 2). Venn diagram was plotted using Venny, an interactive tool for comparing list.

**Figure 4:**
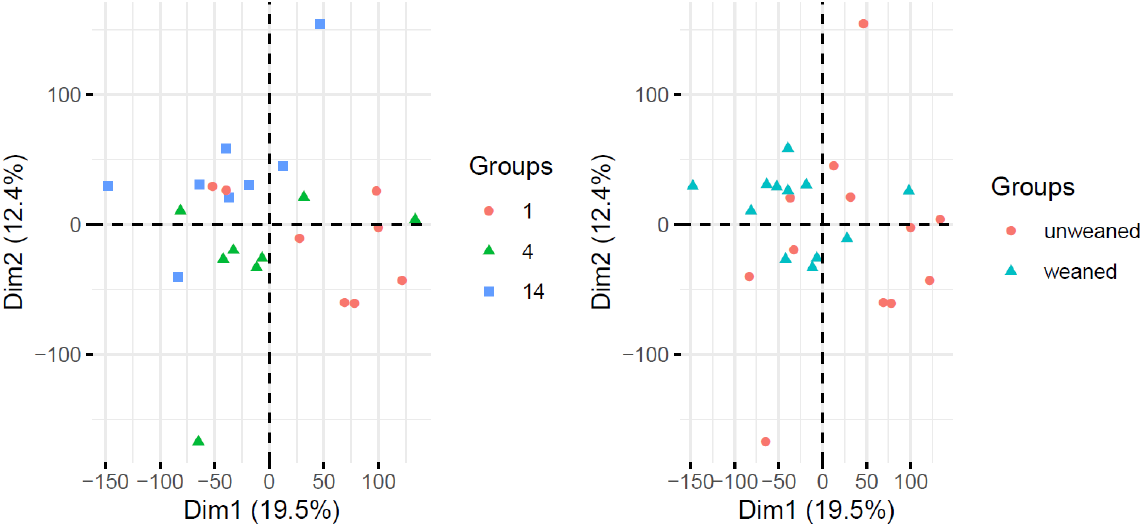
Principal Component Analysis (PCA) of gene expression data from pigs across time point and weaning status.

The full list of differentially expressed transcripts at Qval ≤ 0.1 & FC ≥ 2 is provided in Supplementary Table S1. DETs with FC>2 across all time points were subject to Gene Ontology (GO) and KEGG pathway analysis using NIPA https://github.com/richarddemes/NIPA to identify significantly enriched pathways between weaned and unweaned pigs at Qval ≤ 0.05 and minimum number of gene in term of two (Supplementary Table S2). Top 10 significantly enriched GO terms and KEGG pathways are shown in Figure 5.

**Figure 5:**
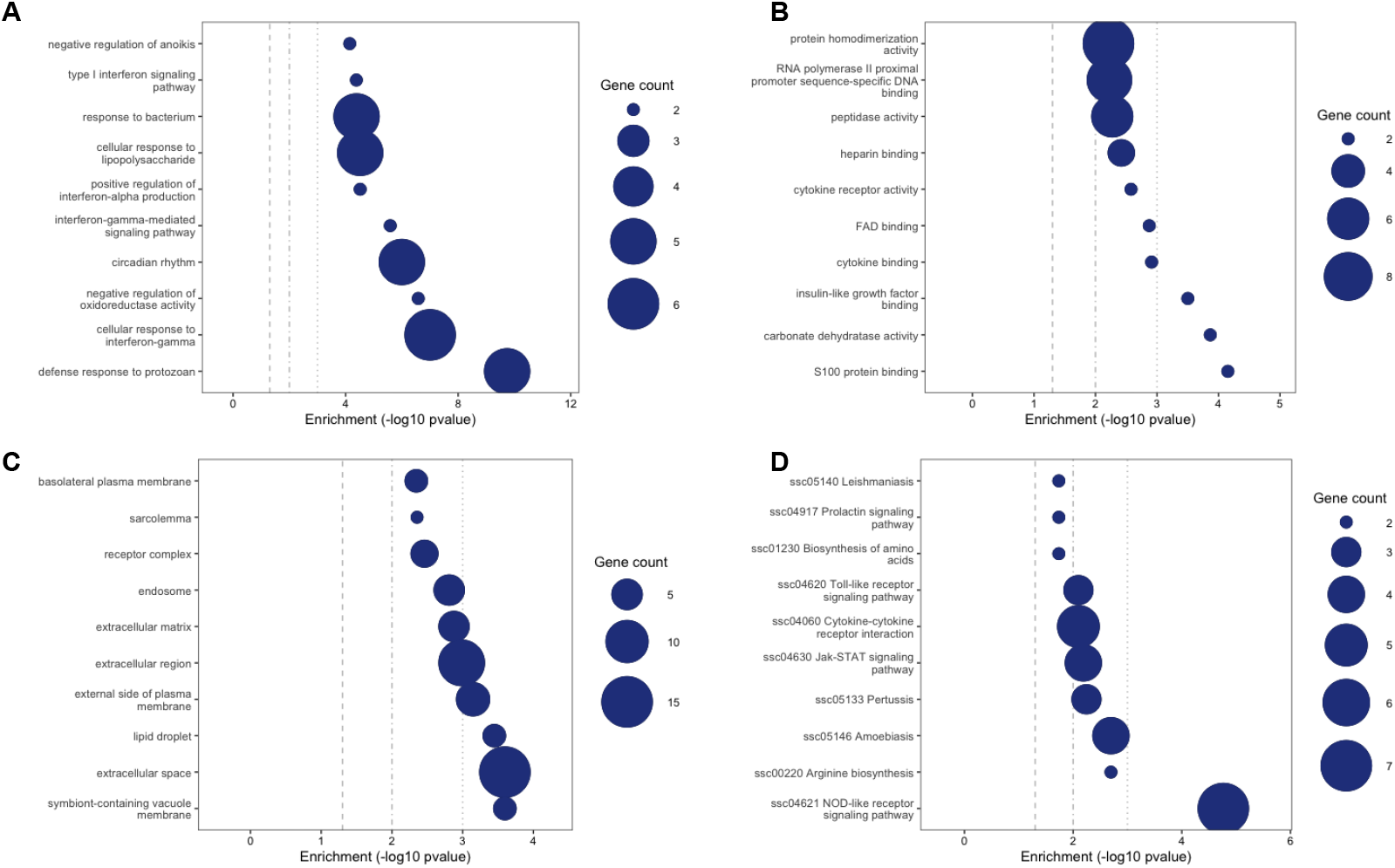
Gene ontology (GO) and KEGG pathway enrichment analysis showing top 10 enriched terms between weaned and unweaned pigs (all time points). Size of circles represents number of gene in each term. A: GO Biological Process, B: GO Molecular Function, C: GO Cellular Compartment, D: KEGG.

## Discussion

In this study, the typical characteristics previously reported in other studies were observed validating our weaning model. Piglet weights were negatively affected by weaning at day 1 and day 4 but recovered at day 14. Post-weaning growth check is a commonly reported problem in pigs with detrimental impact on life-time growth performances representing a large economic loss to the industry (Collins et al., 2017)

Evidence of stress was also observed in this study with elevated plasma cortisol and increased lesion scores in the weaned pigs. Following post-weaning mixing, intense aggressive patterns especially directed to facial, belly and anogenital regions has been reported in previous studies and were shown to be linked with the establishment of social hierarchy between unacquainted pigs (Meese and Ewbank, 1973; Turner et al., 2009). In agreement with previous studies, the majority of lesions found in this study were found on the head, ears and shoulders (data not shown), suggesting aggressive frontal confrontation, however no direct behavioural observation were performed so other behaviour which could have caused lesions could not be excluded. Interestingly, lesion scores in the unweaned piglets were found at 14 d, when pigs were 35 day of age and had reached over 12 kg, suggesting that despite creep feed available, they may have been some possible agonistic behaviour between littermates in competition for food source as the milk yield from the sow starts to decrease at this stage of lactation (Hansen et al., 2012). Weaning age is a controversial topic of discussion regarding performance and welfare, with typical weaning age varying between production systems and countries. Most studies agree that weaning at an early age (<21 days old) exacerbate the effect of weaning stress (Smith et al., 2010; Xun et al., 2018) but our results suggest that later weaning at 35d could also have negative impact on the piglets welfare.

Measurements of intestinal structures such as villus and crypt have been commonly used to evaluate gut health as a measure of absorptive capacities for nutrients (Bontempo et al., 2006; Le Bon et al., 2010). As reported by others, the current study shows that villus morphometry is impaired by weaning. Although feed intake was not measured in this trial, low feed intake post-weaning is considered one of the main aetiological factors for morphological changes in gut physiology due to the importance of enteral stimulation for mucosal homeostasis (Spreeuwenberg et al., 2001). Villus atrophy during the first few days post-weaning has been commonly reported and is often associated with loss in barrier function, decreased enzyme activity and decreased performance such as seen in this study (Lallès, 2008). Interestingly, the effect observed on the crypt depth may indicate an increased level of cellular activity and turnover in the intestinal mucosa in response to weaning.

This study has revealed a large number of genes up and down regulated as consequences of weaning in the pig colon highlighting the profound physiological impact of weaning on healthy pigs. As the genome of the domestic pig is relatively recent in comparison with other model species (Groenen, 2016), many genes and transcripts still have limited annotation and identified function to this date. Differential gene expression in the gut of weaned pigs has previously been investigated using microarray-based techniques (Inoue et al., 2015; Wang et al., 2008), however the number of genes and pathways identified were limited to known transcripts, while RNA seq platform offers a more comprehensive view of the transcriptome including genes of unknown functions. Freeman et al. (2012) documented the first transcriptomic atlas of the domestic pig describing tissue-specific gene expression and emphasised the importance to generate and increase our knowledge of transcriptomic expression to document and understand the function of many genes with limited functional annotation. The transcriptomic profile of pig along the small intestine has been previously evaluated in the duodenum, ileum, jejunum and ileal Peyer’s Patches of healthy growing pigs (Mach et al., 2014), but relatively little is known about the colon in pigs. The current study identified many genes associated with weaning with and without known function, contributing to advancing out fundamental understanding of the pig genome.

The large majority of differentially expressed transcripts were found to peak on day 1 post weaning and progressively returned to unweaned level by day 14, emphasising the abrupt response of pigs to weaning (Lallès et al., 2007). There was little overlap in transcripts between time points suggesting a coordinated regulation of the host in response to weaning. These findings could inform selection of appropriate time points for further trials in weaned pigs.

Due to the large number of DET identified in this study, the main focus of this discussion is on pathways significantly affected by weaning rather that individual genes. Metabolic, immune and barrier function activities have previously been implicated with weaning response in both the small and large intestine but the underlying mechanism for these changes have not been previously identified. In the current study, we found several pathways related to microbial response and immune functions activated post-weaning; specifically, NOD-like receptors, Toll-like receptor and JAK-STAT signalling pathways are shown to be significantly activated. These signalling pathways indicate a cellular response to Microbial Associated Molecular Patterns (MAMPs) that drives the activation innate immune response. As such, we also observe positive regulation of Interferon alpha production and Interferon gamma mediated pathways.

In agreement with these results, Wang et al. (2008) also reported that weaning at 21 days of age showed an increase in expression of genes associated with oxidative stress and immune activation but decreased expression of genes related to nutrient utilisation and cell proliferation in the jejunum of pigs using microarray analysis. A previous study also reported that pigs weaned at a later age (28 days) showed increase in pro-inflammatory cytokines expression in the small intestine and colon during the first two days post weaning before returning to pre-weaned level after 8 days post weaning (Pié et al., 2004). The interferon response following weaning stress in pigs has previously been characterised and showed that weaning causes the release of IFN-α and the transient shut-off of the corresponding gene transcriptions in PBMC (Razzuoli et al., 2011). Bailey (2009) also demonstrated that weaned pigs develop an early and transient immune response to novel dietary antigens at weaning before establishing tolerance (Everaert et al., 2017).

In addition, Moeser et al. (2017) demonstrated that colonic and jejunal transepithelial resistance was increased resulting in impaired permeability of the gut barrier as a result of weaning, which may facilitate infiltration of luminal component such as bacteria or bacterial products. Here we see at least two GO biological terms that would agree with this hypothesis: “cellular responses to LPS” and “response to bacterium” where genes such as NOS2, CD274, IRF3, CXCL11 and ACOD1, are upregulated.

The immune regulation observed through our transcriptomic results also reflects the significant increase in granulocyte levels found in blood, suggesting both a local and systemic activation of immunity in response to weaning. The combined effect of immune and stress activation can lead to an energy cost whereby growing animals divert their energy resource towards these responses instead of growth which can explain reduced performance observed in weaned pigs. In growing pigs, transcriptomic multi-tissue analysis revealed that activated immune response, protein metabolism, defence against pathogens and oxidative stress were the main biological pathways associated with feed efficiencies (Gondret et al., 2017). The authors suggested that dietary intervention with anti-inflammatory or antioxidant properties could be evaluated to improve efficiencies in growing pigs. As feed accounts for more than 60% of the cost of food production, improving feed efficiency is a major target to improve profitability of the pig industry (Jing et al., 2015).

In recent years, significant progress has been made in characterising the complex network of communication and signals between the nervous, immune and endocrine systems of the gut. The role of the gut-brain axis in neurological disorders is well recognised but its mechanism poorly understood. Activation of the gut brain axis under stressful stimuli has been shown to stimulates inflammatory pathways, alteration of gut barrier function and has been associated with changes in intestinal microbiota in humans and animal models (Martin et al., 2018). The multifactorial challenges faced by pigs at weaning have provoked changes in behavioural response and elevated corticoids levels as shown in previous studies (Campbell et al., 2013; Flynn and Wu, 1997; Li et al., 2016). These could be in part responsible for the immune activation observed at the transcriptomic level in the colon but would require further investigation to determine if these changes can be attributed to brain.

The colon also harbours the richest and most diverse microbial population of the gut. In recent years, a large number of studies have documented the development of the gut microbiota of the pig over time (Frese et al., 2015; Guevarra et al., 2018b; Han et al., 2018; Ke et al., 2019; Mach et al., 2015; Wang et al., 2019). As a transition from a milk-based to a plant-based diet, weaning is typically associated with a decrease in gut microbial diversity and major shifts in bacterial taxa composition (Guevarra et al., 2019). In line with other inflammatory gut disorders, such as IBD or Crohn’s disease, the causal relation effect between inflammation and microbiota is unclear. Recently, several studies have provided evidence to suggest that inflammation in gut tissue is conducive to the proliferation of gut bacteria that contribute to pathogenesis or disease development (Zeng et al., 2017). In the current study, we observe activation of immune signalling pathways and pro-inflammatory cytokines in the early days post weaning, and we also observe regulation of pathways involved in oxido-reducatase activity. Amongst the activated pathways related to anti-microbial defence, genes involved in oxidative burst such as NOS2 (Nitric oxide synthesase 2) and NOX1 (NAPDH oxidase 1) are upregulated which are involved in the production of nitic oxide and superoxide radicals. These nitrogen rich compounds are rapidly converted into nitrate (NO3-) in the lumen providing favourable condition for the growth and proliferation of gut bacteria that carry nitrate reductase genes such as *Enterobacteriaceae* including *E. coli* and *Salmonella* (Winter et al., 2013). In addition, inflammatory conditions provide increased level of luminal oxygen due to elevated blood flow and haemoglobin, this favours aerobic respiration of *Enterobacteriaceae* while inhibiting the growth of obligate anaerobes such as *Bacterioides* and *Clostridia* (Zeng et al., 2017). *Enterobacteriaceae* have a detrimental effect on pig health and growth and is one of the leading cause of diarrhoea in pig production (Rhouma et al., 2017). *E. coli* is highly prevalent post-weaning and can lead to mortality and zoonosis (Luppi, 2017). As a result, antibiotic usage is commonly used to treat and prevent *E*.*coli* infection and have led to increasing reports in colistin-resistant *E. coli* found in pigs (Rhouma *et al*., 2017). Tackling post weaning inflammatory response could represent a step towards reduction in antimicrobial use in pig production.

At weaning, pigs are abruptly transitioned from sow’s milk to a complex plant-based diet with distinct nutritional profile and as such transcriptomic modulation of pathways involved in nutritional metabolism would be expected. Whilst the majority of nutrient digestion and absorption takes place in the small intestine, the main role of the colon in digestive functions is to reabsorb water and electrolytes. However, in the current study, biosynthesis of amino acids, in particular arginine biosynthesis was the second most significant KEGG pathway enriched in weaned pigs via the up regulation of argininosuccinate synthase 1 gene (ASS1). Arginine is an essential amino acid that play key roles in nutrition and metabolism. Young mammals, including piglets, have a particularly high requirement of arginine for growth and metabolic function. As enterocytes actively transport and metabolise arginine, the gut is an important organ for maintaining body arginine homoeostasis (Wu et al., 2018). L-Arginine is also the biological precursor of nitric oxide (NO), and alteration of arginine uptake and metabolism has been found to be associated with inflammatory bowel diseases, such as ulcerative colitis (Coburn et al., 2016; Luiking et al., 2012; Stuehr, 2004). In post-weaning pigs, arginine metabolism in the gut is critical to maintain normal intestinal physiology and for efficient utilisation of dietary protein (Wu et al., 2004). In addition, glucocorticoids plays an important role in regulating the enhanced arginine metabolism in the enterocytes of post-weaning pigs (Flynn and Wu, 1997). Arginine availability in the digestive tract plays a key role in maintaining intestinal immune homeostasis under conditions of inflammation and infection (Das et al., 2010; Fritz, 2013; Singh et al., 2019). A number of studies have reported that arginine administration significantly attenuate intestinal inflammation under physiological and pathological conditions including intestinal dysfunction in weaned piglets, and enhancement of growth performance and survival (Che et al., 2019; Wu et al., 2010; Zheng et al., 2018). Other models have reported that arginine supplementation also down-regulated JAK-STAT signalling pathway and attenuated the inflammatory response, which exerted protective effects on the intestine of chickens challenged with *C. perfringens* ((Zhang et al., 2019). Arginine has been shown to exert anti-inflammatory and antioxidant effects in IPEC-J2 cells challenged with LPS (Qiu *et al*., 2019). Reports of reduced arginine availability in conditions of acute and chronic stress, often associated with increase in NOS2 activity are in agreement with the results of the current study (Luiking et al., 2012). The specific mechanisms of regulation and interaction between cortisol, NOS, immune response, and arginine metabolism observed in this study remain unknown, but could provide further evidence to suggest that arginine requirement should be carefully evaluated when designing diet to support pigs during the weaning transition.

Finally, although the current study has identified a large number of transcripts and pathways regulated at the mRNA level, it is likely that post-transcriptional and post translational regulatory mechanisms also regulate the host response to weaning and should be investigated in future studies as comprehensive and complementary approach to transcriptomic methods.

## Conclusion

Weaning is a multifactorial event that results in complex interactions between gut, brain and metabolism. Understanding these responses and the molecular mechanisms that underpins these changes is critical to improve sustainable pig production. This study has identified multiple genes and pathways differentially regulated by weaning. These results revealed that pigs going through the weaning transition undergo a transient period of inflammatory state with temporary breakdown of barrier functions in the gut. The condition of the inflamed gut have been previously shown to provide favourable growth advantage for the expansion of *Enterobacteriaceae*, a leading cause of enteric disease in pigs. Under the experimental and controlled conditions of this trial, differential gene expression returned to unweaned control levels by day 14 post weaning. However, the translation of the study results to commercial production setting remains to be explored.

Indicators of weaning stress and response have previously been used including histology, systemic markers of immunity and characterisation of the microbiota composition. Here, we have identified a number of target gene and pathways that could also be used as biomarker of intestinal inflammation to complement these measures. Together, these could provide valuable tools to monitor host response post-weaning, especially in context of intervention strategies aimed to reduce antibiotic use and improve pig health and performance. Finally, as weaning in pigs have been used as a model for stress-related bowel dysfunction in humans, it would be of interest to investigate if the similar transcriptomic changes are involved in these disorders.

## Acknowledgements

We acknowledge Lallemand Inc, the University of Nottingham and Nottingham Trent University for funding. We also acknowledge the team at Bio-Support Unit at the University of Nottingham for help in running the trial. We thank Prof. Ian Connerton for critical reading and review of the manuscript.

